# Mirage: A phylogenetic mixture model to reconstruct gene-content evolutionary history using a realistic evolutionary rate model

**DOI:** 10.1101/2020.10.09.333286

**Authors:** Tsukasa Fukunaga, Wataru Iwasaki

**Affiliations:** Department of Computer Science, Graduate School of Information Science and Technology, The University of Tokyo, Tokyo, Japan; Department of Biological Sciences, Graduate School of Science, The University of Tokyo, Tokyo, Japan; Department of Computational Biology and Medical Science, Graduate School of Frontier Sciences, The University of Tokyo, Chiba, Japan; Atmosphere and Ocean Research Institute, The University of Tokyo, Chiba, Japan; Institute for Quantitative Bioscience, The University of Tokyo, Tokyo, Japan; Collaborative Research Institute for Innovative Microbiology, The University of Tokyo, Chiba, Japan

**Keywords:** Genome evolution, Gene content, Probabilistic evolutionary model, Phylogenetic mixture model, Maximum likelihood method

## Abstract

Reconstruction of gene-content evolutionary history is an essential approach for understanding how complex biological systems have been organized. However, the existing gene-content evolutionary models cannot formulate complex and heterogeneous gene gain/loss processes, which reflect diverse evolutionary events and greatly depend on gene families. In this study, we developed Mirage (<u>MI</u>xture model with a <u>R</u>ealistic evolutionary rate model for <u>A</u>ncestral <u>G</u>enome <u>E</u>stimation), which allows different gene families to have flexible gene gain/loss rates, but reasonably limits the number of parameters to be estimated by the expectation-maximization algorithm. Simulation analysis showed that Mirage can accurately estimate complex and heterogeneous gene gain/loss rates and reconstruct gene-content evolutionary history. Application to empirical datasets demonstrated that our evolutionary model better fits genome data from various taxonomic groups than other models. Using Mirage, we revealed that gene families of metabolic function-related gene families displayed frequent gene gains and losses in all taxa investigated. The source code of Mirage is freely available at https://github.com/fukunagatsu/Mirage.

## 1. Introduction

Gene gain and loss events in genomes have played essential roles in the evolutionary history of life. Complex biological systems that function through the coordination of numerous genes, e.g., metabolic pathways and signal transduction systems, have been constructed through the accumulation of such events. To answer the fundamental biological question of how such complex systems have been organized, reconstruction of gene content evolutionary history has been established as an important field in bioinformatics (Iwasaki and Takagi 2009; Montague *et al*. 2014; Fernández and Gabaldón 2020). To date, various algorithms to reconstruct gene content evolutionary history have been developed (Snel *et al*. 2002; Hahn *et al*. 2005; Iwasaki and Takagi 2007; Csurös and Miklós 2009; Cohen and Pupko 2010; Ames *et al*. 2012; Li *et al*. 2014; Zamani-Dahaj *et al*. 2016; Li *et al*. 2019). These algorithms estimate the gene content of ancestral species based on the maximum parsimony (MP) or maximum likelihood (ML) method, where the ML method is known to show better performance (Cohen and Pupko 2011; Ames *et al*. 2012).

In the ML method, it is important which gene-content evolutionary model is adopted. Given an ortholog table and a phylogenetic tree, the ML method first estimates the evolutionary model parameters, such as gene gain and loss rates, using numerical optimization methods or the expectation-maximization (EM) algorithm. Then, the ML evolutionary history of gene-content is reconstructed based on the estimated parameters. Some ML methods adopt a two-state evolutionary model and require a two-state ortholog table, which contains the presence/absence information of each ortholog group in each genome, and estimate whether each ortholog group existed or not at each ancestral node of the given phylogenetic tree (Cohen and Pupko 2010; Li *et al*. 2014) (Fig. 1A). Although the two-state evolutionary model is mathematically simple, it is apparently unable to deal with gene copy number variations, which play important roles in the evolution of biological systems (Saitou and Gokcumen 2020). At the opposite extreme, the ML method can also adopt an “infinite-state” (or least-constraint) evolutionary model and require a “raw” ortholog table, which contains copy number information of each ortholog group in each genome, and estimate the copy number of each ortholog group at each ancestral node (the least-constraint model; Fig. 1B). A major drawback of this approach is that parameter numbers can become too large to be estimated. For example, even if the upper bound of the gene copy numbers is set to ~1,000 (cf., olfactory receptor genes (Niimura 2009)), ~2,000 parameters still need to be estimated at a huge computational cost and overfitting risk.

**Fig. 1.**
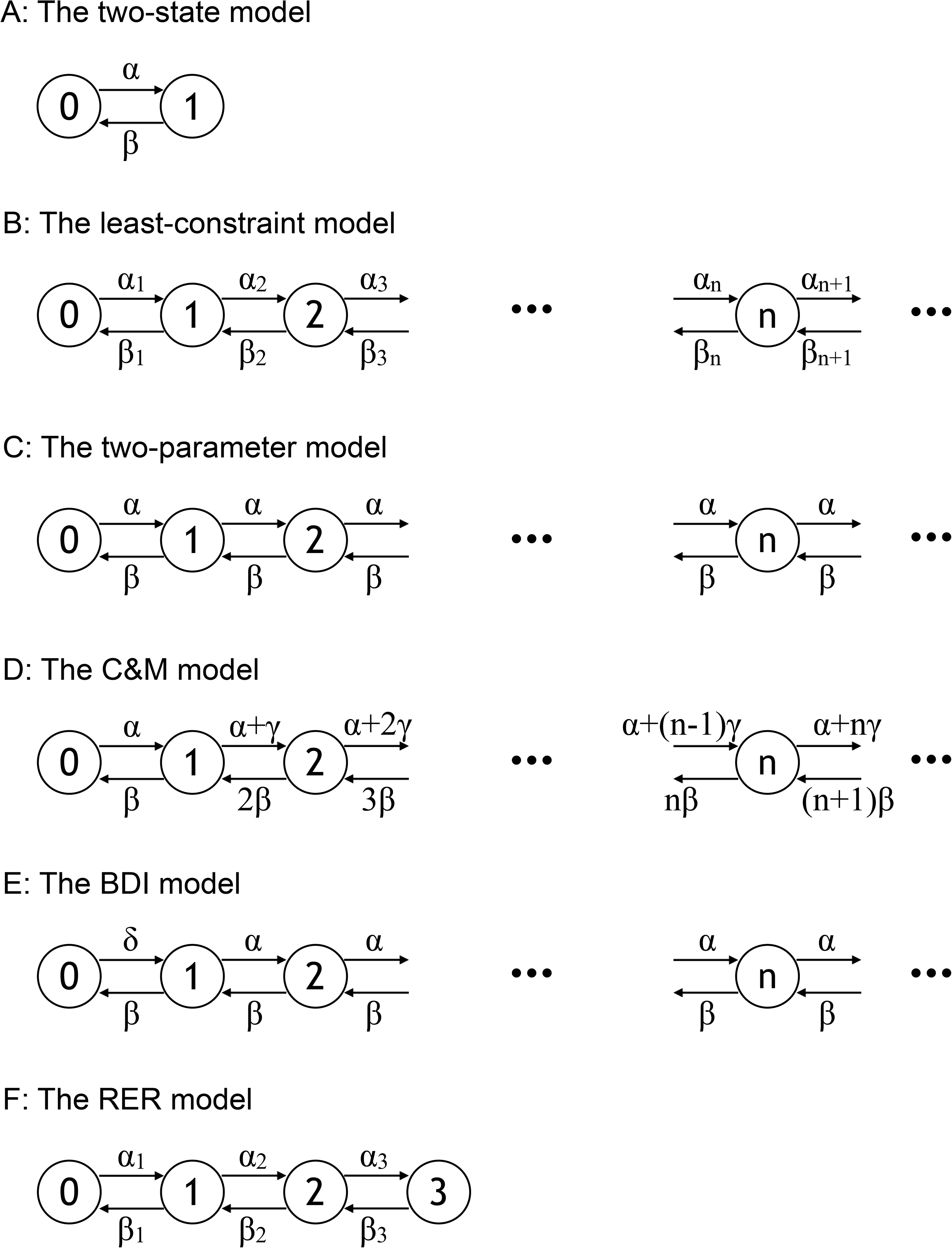
Schematic illustration of the evolutionary models. Enclosed numerals and arrows indicate gene copy numbers and the gene gain/loss events, respectively. (A) Two-state, (B) least-constraint, (C) two-parameter, (D) C&M, (E) BDI, and (F) RER evolutionary models are shown.

Between the two extremes of the two-state and least-constraint models, previous studies on the ML method have adopted gene-content evolution models that consider gene copy numbers but limit the numbers of parameters. The simplest model is the two-parameter model, which considers only gene gain and loss rates (Fig. 1C) (Hahn *et al*. 2005; Iwasaki and Takagi 2007). Other models (Csurös and Miklós (C&M) model and Birth, Death and Innovation (BDI) model) decompose the gene gain rate into gene birth (innovation) and duplication rates, resulting in three parameters (Figs. 1D and 1E) (Csurös and Miklós 2009, Karev *et al*. 2002; Ames *et al*. 2012). However, these two- and three-parameter models still cannot formulate complex processes that reflect diverse evolutionary events, e.g., *de novo* gene birth, tandem duplication, ectopic recombination, and horizontal gene transfers (HGTs). These models ignore tendencies that gene gain rates change non-linearly in accordance with gene copy numbers in case of tandem duplications, and copy numbers of many genes are kept one because of dosage balance conservation.

Another aspect of gene-content evolution that should be considered is that different gene families have different gene gain/loss rates. For example, housekeeping genes are seldom lost from genomes and gene loss rates to zero copies are small for such genes, whereas antibiotic genes are easily lost from genomes. Another example is olfactory receptor genes, which are prone to increase copy numbers and have exceptionally large gene gain rates. To deal with such heterogeneity in gene-content evolution, the phylogenetic mixture model, which is widely used in molecular evolutionary analyses to handle heterogeneous evolutionary rates among different residues (Lartillot and Philippe 2004; Pagel and Maede 2004; Quang *et al*. 2008; Dang and Kishino 2019), can be adopted. However, in gene-content evolutionary analysis, the phylogenetic mixture model has been adopted only in the ML-based methods that use the two-state model (Spencer and Sangaralingam 2009; Cohen and Pupko 2010; Li *et al*. 2014; Zamani-Dahaj SA *et al*. 2016).

In this study, we proposed a realistic evolutionary rate (RER) model, a novel gene-content evolutionary model, and developed Mirage (<u>MI</u>xture model with a <u>R</u>ER model for <u>A</u>ncestral <u>G</u>enome <u>E</u>stimation), which adopts the RER model and phylogenetic mixture model for accurate ML reconstruction of gene-content evolutionary history (Fig. 1F). To limit the number of model parameters to be estimated, the RER model enables users to set an upper bound of gene copy numbers (Iwasaki and Takagi 2007; Ames *et al*. 2012). We verified that Mirage can estimate both model parameters and gene-content evolutionary history with high accuracy using simulated datasets. In addition, we demonstrated that the combination of the RER model and phylogenetic mixture model fitted empirical datasets better than the other models. Finally, we reconstructed gene-content evolutionary histories of several taxonomic groups using Mirage, and revealed that gene families involved in metabolic functions frequently exhibited gene gain/loss events in all taxonomic groups investigated.

## 2. New Approaches

### 2.1 Input data

The input data for our method are an ortholog table *D* and a phylogenetic tree *T*. *D* is a data matrix that consists of *N* species (genomes) and *L* gene families (ortholog groups). *D*_*i*,*j*_, which is an element of the species *i* and the gene family *j* in the matrix, represents the gene copy number of *j* in *i*. The phylogenetic tree *T* is a binary rooted tree whose branches have branch lengths greater than 0. The tree has *N* leaves (external nodes), which correspond to the *N* species in the ortholog table *D*. *T* also has *N* − 1 internal nodes, which correspond to the ancestral species. The reconstruction problem of gene content evolutionary history is defined as an estimation problem of the gene copy numbers in the ancestral species for each gene family (*X*).

### 2.2 The RER model and the phylogenetic mixture model

Gene-content evolution is formulated as a continuous-time Markov model, where gene copy numbers and gene gain/loss events are represented as states and state transitions, respectively. Gene gain/loss events in each gene family are assumed to have occurred independently of those in other gene families. In an infinitesimal time *Δt*, a gene gain/loss event of one gene is assumed to have occurred at most once. In the two-parameter model, where the model parameters are a gene gain rate α, and a gene loss rate *β*, transition from a gene copy number *n* to *n* + 1 and *n* − 1 occur at probabilities of, *αΔt* and *βΔt*, respectively, in *Δt* (Fig. 1B) (Hahn *et al*. 2005; Iwasaki and Takagi 2007). Csurös and Miklós developed a model with three parameters: a gene acquisition rate α, a gene loss rate *β* and a gene duplication rate 0 (Fig. 1C) (Csurös and Miklós 2009). By assuming that HGT is a main mechanism of gene acquisition, the C&M model defines transition rates from *n* to *n* + 1 and *n* − 1 as α + *nγ* and *nβ*, respectively (note that the gene gain/loss rates change linearly with gene copy numbers). The other three-parameter model, the BDI model, utilizes a novel gene family acquisition rate δ in addition to a general gene gain rate α and a gene loss rate *β* (Fig. 1D) (Karev *et al*. 2002; Ames *et al*. 2012). This model is basically the same as the two-parameter model, except that the transition rate from 0 to 1 is δ.

In this study, we proposed a RER model, which allows all state transition rates to be different. Instead, to limit parameter numbers, we set the maximum gene copy numbers as a parameter *l*_*max*_ (i.e., gene families having copy numbers larger than *l*_*max*_ are considered to have *l*_*max*_ gene copies) (Fig. 1E). That is, the number of states is *l*_*max*_ + 1 (0 to *l*_*max*_). Because *l*_*max*_ is a user-input parameter, the user can freely set it to a reasonable value according their interest (for example, if the user is interested in the evolution of many copy gene families, *l*_*max*_ can be set large at the expense of a parameter number increase). Let *R* be a (*l*_*max*_ + 1) × (*l*_*max*_ + 1) transition rate matrix. [*R*]_*i*,*j*_, which is an (*i,j*)-th element of *R*, represents the state transition rate from the state *i* to the state *j* in *Δt*. *[R]*_*i,j*_ = 0 when |*i* − *j*| > 1, and the number of evolutionary rate parameters is 2*l*_*max*_. We define *p*(*y*|*x*, *R*,*t*) as a transition probability from state *x* to *y* in time *t*. If *t* = 0, the gene copy number does not change and *p*(*y*|*x*, *R*,0) = [*I*]_*x*,*y*_, where *I* is the identity matrix. If *t* = *Δt*, *P*(*y*|*x*,*R*,*Δt*) = [*I* + *RΔt*]_*x,y*_, where [*R*]_*i*, *j*_ = −Σ_*j*,*i*≠*j*_[R]_*i,j*_. Then, under the Markov process assumption, we obtain 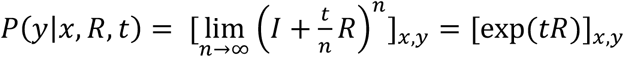.

Furthermore, to allow different gene families to have different gene gain/loss rates, we combined the RER model with the phylogenetic mixture model. Instead of assuming that all of the *L* gene families evolve under the same transition rate matrix *R*, the phylogenetic mixture model introduces *K* transition rate matrices (i.e., *R*_1_,…, *R*_*K*_). Each of the *L* gene families is probabilistically assigned to *K* clusters as a mixture model. Here, *K* is a user-input parameter.

### 2.3 Parameter estimation and gene-content evolutionary history reconstruction algorithm

The evolutionary model parameter to be estimated is *θ* = {*φ*, *R*_1_,…, *R*_*K*_, *π*_1_,…, *π*_*K*_}, where *φ* is a *K*-length vector that is the mixing probability of each gene-content cluster, *R*_*i*_ is a transition rate matrix of the *i*-th gene-content cluster, and *π*_*i*_ is a (*l*_*max*_ + 1)-length vector that is the state occurrence probability at the root node in the phylogenetic tree of the *i*-th gene-content cluster. Note that the degree of freedom is 3*l*_*max*_*K* + *K* − 1.

The model parameters are estimated by the EM algorithm (Dempster *et al*. 1977). The EM algorithm is an ML method for estimating parameters from observed data in statistical models that assume unobserved hidden states. In our model, the observed data is the ortholog table *D*, while the unobserved hidden states are the gene-content evolutionary history *X* and assignments of each gene family to each gene-content cluster *Z*. The EM algorithm consists of the following four steps. (1) Initialize the model parameter *θ*_*old*_ randomly. (2) Calculate *p*(*X,Z*|*D*, *θ*_*old*_). (3) Calculate *θ*_*new*_ = argmax_*θ*_*Q*(*θ*,*θ*_*old*_) where *Q*(*θ*, *θ*_*old*_) = *Σ*_*X,Z*_ *p*(*X,Z*|*D*, *θ*|*θ*_*old*_)In*p*(*X,Y,Z*|*θ*). (4) If the log-likelihood converges, terminate the EM algorithm. Otherwise, substitute *θ*_*new*_ for *θ*_*old*_ and return to the step (2). The steps (2) and (3) can be efficiently calculated using the eigenvalue decomposition of the state transition probability matrices and a dynamic programming method across the phylogenetic tree *T* (Holmes and Rubin 2002; Quang *et al*. 2008; Kiryu 2011).

After the parameter estimation, the ML evolutionary history 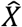 is reconstructed by a dynamic programming method using the estimated parameters. The reconstruction method is similar to the Viterbi algorithm, which obtains the ML path of hidden states in the hidden Markov model, and also resembles an algorithm for the reconstruction of ancestral protein sequences (Pupko *et al*. 2000).

The details of the algorithms are described in the Supplementary Materials. We implemented the algorithms in C++. The source code is freely available at https://github.com/fukunagatsu/Mirage.

## 3. Results

### 3.1 Performance evaluation of Mirage based on simulated datasets

We first evaluated the performance of Mirage using simulated datasets. We simulated the evolution of 10,000 gene families in a phylogenetic tree that had 128 leaves for 10 times, creating 10 datasets. We used the RER model whose maximum gene copy number *l*_*max*_ was set to 3 and the phylogenetic mixture model whose number of gene content cluster *K* was set to 4 for the simulation. Details are described in the Material and Methods.

Then, we applied Mirage to the 10 simulated datasets to estimate the evolutionary model parameters and reconstruct the gene-content evolutionary history. Fig. 2 shows the relative errors of the estimated model parameters. The medians and maximum values of the relative errors were 1.25 and 7.23 for *φ*, 1.122 and 27.164 for *R*, and 14.7565 and 510.67 for *π*, respectively. These results imply that Mirage can estimate parameters with high accuracy, although the estimated *π* may substantially differ from the true parameter. Such difficulty in root parameter estimation is well known and may stem from the fact that the root node is the topologically the furthest away from the observable leaf nodes (i.e., extant genomes).

**Fig. 2.**
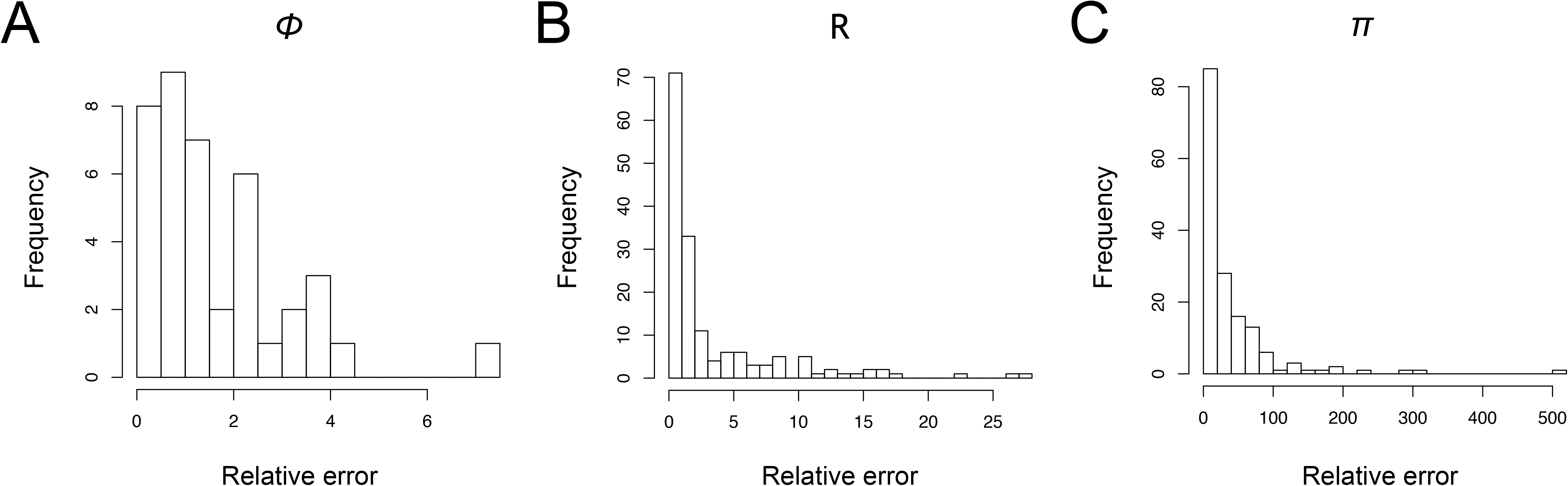
Distributions of the relative errors of the estimated parameters for the simulated datasets. A relative error is 100 × |the estimated value − the true value|/the true value. The x-axis and y-axis represent the relative error and the frequency, respectively. Distributions for (A) φ, (B) R, and (C) *π* are shown. Note that R and *π* have many parameters and thus their frequencies are larger than that of φ.

The accuracy of the reconstructed ancestral states (gene copy numbers) was 84.0% (Table 1). When the phylogenetic mixture model was not used (i.e., *K* = 1), the accuracy decreased to 81.6%, likely because the heterogeneity of gene-content evolution was ignored. If the copy number states were ignored and only presence/absence information was considered (i.e., if incorrect estimation among the copy numbers 1, 2, and 3 was ignored), the accuracy of presence/absence state reconstruction of the ancestral nodes was 92.0%. On the other hand, when gene copy number information was not used and the two-state model (with the phylogenetic mixture model) was applied (i.e., *l*_*max*_ = 1), the accuracy was 91.9%. This result indicates that the two-state model is sufficiently accurate only to reconstruct presence/absence information. In addition, if the two-parameter, C&M, and BDI models (without the phylogenetic mixture model) were used, the accuracies were 82.8%, 82.6%, and 82.8%, respectively. Note that *l*_*max*_ was set to 3 to compare the models under the same condition.

**Table 1.** Estimation accuracy of the reconstructed gene content evolutionary history

| Method | Estimation accuracy | Estimation accuracy of presence/absence only |
| --- | --- | --- |
| RER, $K = 4$, $l_{max} = 3$<br>(True condition) | **84.0%** | **92.0%** |
| RER, $K = 1$, $l_{max} = 3$ | 81.6% | - |
| RER, $K = 4$, $l_{max} = 1$<br>(Two-state model with<br>phylogenetic mixture model) | - | 91.9% |
| Two-parameter, $K = 1$, $l_{max} = 3$ | 82.8% | - |
| Csurös, $K = 1$, $l_{max} = 3$ | 82.6% | - |
| BDI, $K = 1$, $l_{max} = 3$ | 82.8% | - |

### 3.2 Investigation of gene copy numbers in empirical datasets

We next applied Mirage to empirical datasets. We created three empirical datasets including Archaea (domain), Micrococcales (order), and Fungi (kingdom). We used ortholog tables provided in the STRING database (Szklarczyk *et al*. 2019). We downloaded phylogenetic trees of Archaea and Micrococcales from the Genome Taxonomy Database (Parks *et al*. 2019) and one of Fungi from the SILVA database (Yilmaz *et al*. 2014; Yarza *et al*. 2017). The Archaea, Micrococcales, and Fungi datasets comprised 167 species and 11,717 gene families, 111 species and 6,653 gene families, and 123 species and 23,791 gene families, respectively.

The largest gene copy numbers among all gene families were 183, 102, and 513 in the Archaea, Micrococcales, and Fungi datasets, respectively, indicating that it would be reasonable to set an upper bound of gene copy numbers to limiting numbers of model parameters. Fig. 3 shows cumulative relative frequency curves of the largest gene copy numbers in each gene family. Approximately 60% of the gene families had one copy at most in any dataset, indicating that the two-state model was not suitable for reconstructing the evolution of approximately 40% of the gene families. In contrast, when we set *l*_*max*_ to 3, the evolution of approximately 90% of the gene families could be reconstructed without the gene copy number cut-off.

**Fig. 3.**
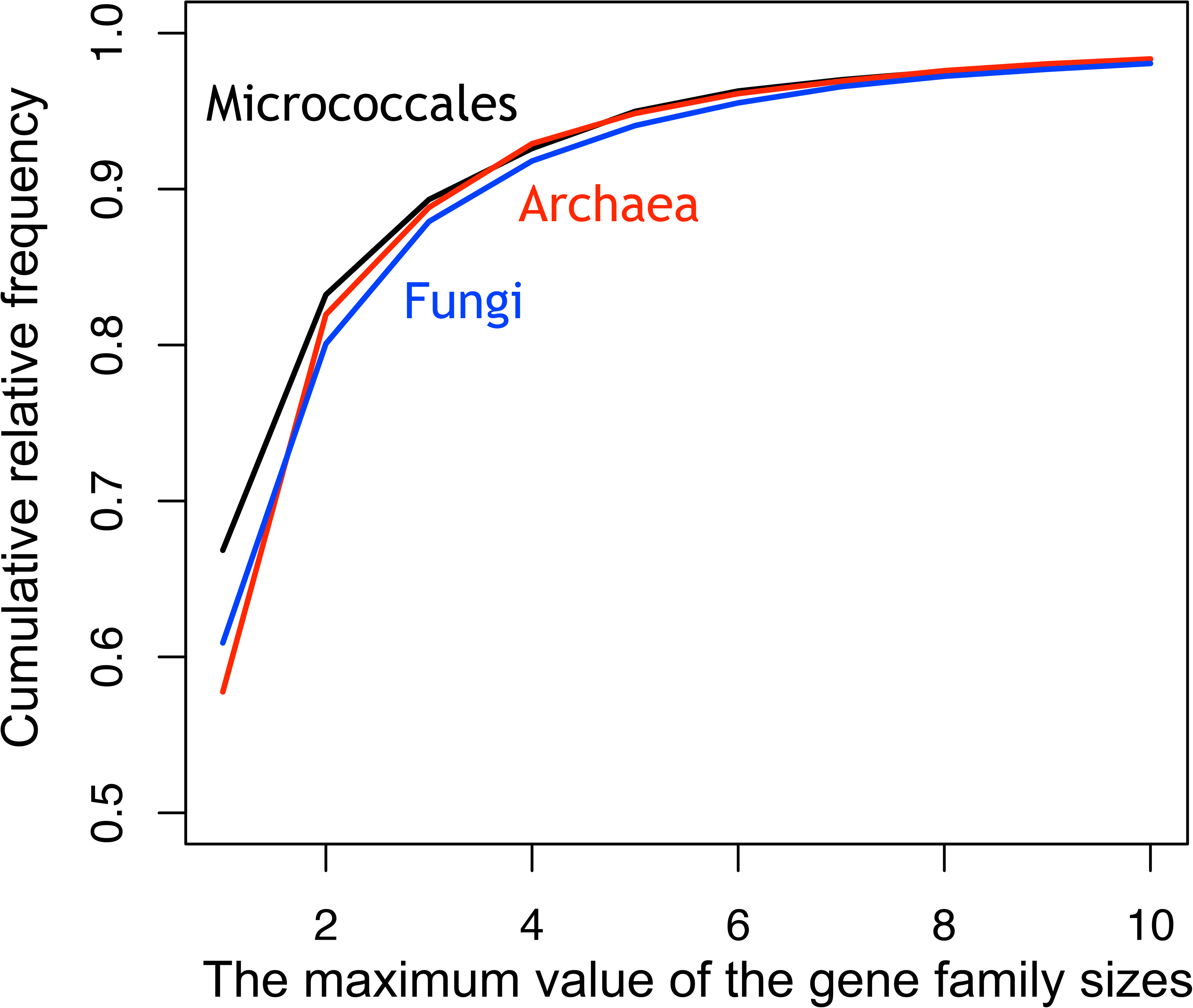
Cumulative distribution curves of the maximum gene copy numbers in the empirical datasets. The x-axis and y-axis represent the maximum value of the gene copy numbers and the cumulative relative frequency, respectively. The Archaea, Micrococcales, and Fungi datasets are represented by red, black, and blue lines, respectively.

### 3.3 Comparison of models and parameters by holdout validation

Next, we compared effects of models and parameters by evaluating holdout performance of Mirage using empirical datasets. We first divided each dataset into gene families of training and test datasets, and estimated the model parameters using the training dataset only. We then calculated log-likelihood of the test dataset. Similarly, parameters were estimated and log-likelihood values were obtained using the two-parameter, C&M, and BDI models for each dataset. Note that *l*_*max*_ was set to 2, 3, or 4 to compare the models under the same condition.

Regardless of *l*_*max*_ or the dataset used, the log-likelihood increased with the increasing number of transition-rate matrices *K*, except for limited cases, likely because of convergence to local optima by the EM algorithm, and the RER and two-parameter models showed the best and the worst log-likelihood, respectively (Fig. 4). When we divided the training and test datasets in different ways (i.e., by species or by species and gene families), we obtained similar results (Figs. S1-S2). In conclusion, both the RER model and the phylogenetic mixture model yielded a gene-content evolutionary model with high log-likelihood values.

**Fig. 4.**
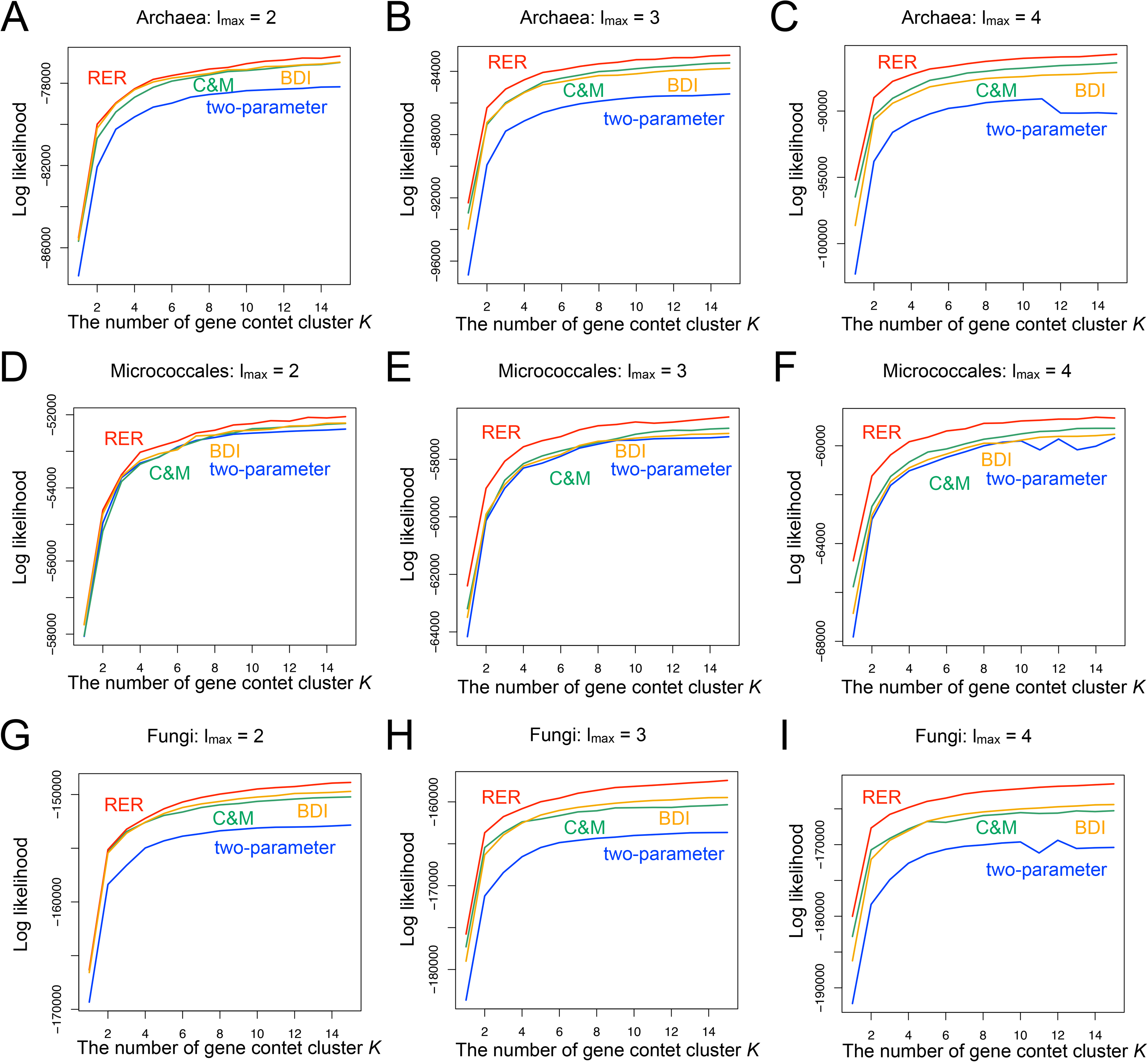
Log-likelihood values of various model settings by the holdout validation of the experiment 1. The x-axis and y-axis represent the number of gene-content clusters L and the log-likelihood of the test dataset, respectively. The two-parameter, C&M, BDI, and RER models are represented by blue, green, yellow, and red lines, respectively. (A-C) Archaea dataset when *l*_*max*_ was set to 2-4, (D-F) Micrococcales dataset when *l*_*max*_ was set to 2-4, and (G-I) Fungi dataset when *l*_*max*_ was set to 2-4.

Interestingly, although the C&M and BDI models had the same number of parameters, their log-likelihood values were slightly different (Fig. 4). When the Archaea or Micrococcales dataset was used and *l*_*max*_ was 3 or 4, the C&M model exhibited a larger log-likelihood. On the other hand, when the Fungi dataset was used, the BDI model exhibited a larger log-likelihood. When *l*_*max*_ ≥ 3, the C&M model naturally assumes that gene duplication and loss rates change linearly with gene copies, whereas the BDI model assumes that gene duplication and loss occur at a constant rate regardless of gene copy numbers (Fig. 1). Thus, the difference likely reflects the nature of gene duplications and losses in prokaryotic and eukaryotic genomes. Specifically, the BDI model may be more suitable for eukaryotic evolutionary processes in which meiotic recombination introduces tandem gene duplications and losses, which are basically independent of gene-copy numbers (Fitzpatrick 2012).

### 3.4 Analysis of estimated evolutionary model parameters

Next, we applied Mirage to each of the complete Archaea, Micrococcales, and Fungi datasets. Based on an investigation of gene copy numbers and holdout validation results, we used *K* = 5 and *l*_*max*_ = Estimated model parameters are presented in Table 2, Fig. S3, and the Supplementary Data. In all datasets, the gene gain rates ([*R*_*K*_]_*i*,*i*+1_) tended to be smaller than the gene loss rates ([*R*_*K*_]_*i*,*i*−1_), being consistent to a previous study (Cohen and Pupko 2010).

**Table 2.** Estimated parameters based on the complete Archaea dataset (see Supplementary Materials for Micrococcales and Fungi datasets)

| Cluster ID | $\varphi$ | $\pi$ | $R$ |
| --- | --- | --- | --- |
| 1 | 0.316 | (0.460, 0.070, 0.021, 0.449)<sup>T</sup> | $\begin{pmatrix} -0.160 & 0.160 & 0 & 0 \\ 7.959 & -9.503 & 1.544 & 0 \\ 0 & 11.795 & -15.237 & 3.442 \\ 0 & 0 & 6.615 & -6.614 \end{pmatrix}$ |
| 2 | 0.306 | $(0.000, 0.951, 0.000, 0.048)^T$ | $\begin{pmatrix} -0.016 & 0.016 & 0 & 0 \\ 2.395 & -2.718 & 0.323 & 0 \\ 0 & 4.553 & -5.570 & 1.017 \\ 0 & 0 & 3.479 & -3.479 \end{pmatrix}$ |
| 3 | 0.291 | $(0.864, 0.115, 0.017, 0.003)^T$ | $\begin{pmatrix} -0.031 & 0.031 & 0 & 0 \\ 0.196 & -0.272 & 0.076 & 0 \\ 0 & 0.822 & -1.125 & 0.303 \\ 0 & 0 & 0.500 & -0.500 \end{pmatrix}$ |
| 4 | 0.049 | $(0.438, 0.414, 0.000, 0.148)^T$ | $\begin{pmatrix} -0.281 & 0.281 & 0 & 0 \\ 0.542 & -1.002 & 0.460 & 0 \\ 0 & 2.190 & -3.203 & 1.013 \\ 0 & 0 & 2.433 & -2.433 \end{pmatrix}$ |
| 5 | 0.038 | $(0.553, 0.070, 0.043, 0.335)^T$ | $\begin{pmatrix} -0.615 & 0.615 & 0 & 0 \\ 2.808 & -4.966 & 2.158 & 0 \\ 0 & 6.036 & -7.925 & 1.869 \\ 0 & 0 & 1.651 & -1.651 \end{pmatrix}$ |

We next examined the evolutionary model parameters estimated for each gene-content cluster and each dataset. To quantify the frequency of gene gain/loss events occur in each cluster *k*, we calculated a normalized cluster evolutionary rate, 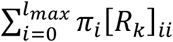 which was divided by the minimum of these values among each dataset (Fig. S4). The maximum normalized cluster evolutionary rates were 60.0, 4.9, and 12.1 for the Archaea, Micrococcales, and Fungi datasets, respectively, indicating that different gene-content clusters have different evolutionary rates. The Archaea dataset exhibited the largest difference, where the gene-content clusters 1, 2, and 3 exhibited large, moderate, and small normalized cluster evolutionary rates, respectively (Table 2). We also investigated whether specific gene functions were enriched in specific gene-content clusters. We used EGGNOG database version 4.0 for gene annotation to COG, arCOG, and NOG gene families and version 3.0 for gene annotation to KOG category (Powell *et al*. 2012; Powell *et al*. 2014). After removing “poorly characterized” supercategories, we observed differences in the enriched COG supercategories among gene-content clusters (Tables S1-S3).

### 3.5 Reconstruction of the gene-content evolutionary history

We then reconstructed the gene-content evolutionary history for each dataset using Mirage. We first counted gene gain/loss events in each gene family from the reconstructed evolutionary history (Fig. S5). In all datasets, gene gain/loss events were rare in most gene families, whereas some gene families exceptionally frequently experienced gene/gain loss events. Tables S4-S6 list the 20 gene families with the most frequent gene gain/loss events for each dataset. Many transposase genes were commonly found in all three datasets, whereas two gene families, COG0286 (HsdM) and COG1848, commonly appeared in the lists of the Archaea and Micrococcales datasets. COG1848 is a function-unknown gene family, whereas COG0286 is annotated as a DNA methylase subunit of the type I restriction-modification system. It is reasonable that this restriction-modification system has been spread by HGT, as is well known for the type II system (Jeltsch and Pingoud, 1996).

Finally, we investigated whether specific gene functions were enriched in the gene families with frequent gene gain/loss events. First, we examined differences in the distributions of the COG supercategories between the top 10% of gene families with frequent gene gain/loss events and entire gene families (Table 3). Based on a chi-square test with Bonferroni’s multiple correction, we found that the “metabolism” supercategory was significantly enriched in the gene families with frequent gene gain/loss events in all datasets. When we analyzed which categories in the “metabolism” supercategory were enriched in the different datasets, we found that different categories were enriched (Table 4). In the Archaea dataset, gene families in categories C, “Energy production and conversion”, and P, “Inorganic ion transport and metabolism”, were the most enriched, probably reflecting the diverse ways in which Archaea obtain energy. In the Micrococcales and Fungi datasets, gene families in categories G, “Carbohydrate transport and metabolism”, and Q, “Secondary metabolites biosynthesis, transport, and catabolism”, were highly enriched, probably reflecting rich secondary metabolism functions of those taxonomic groups.

**Table 3.** Enrichment of COG supercategories by frequent gene gain/loss events.

| COG supercategory | Cellular process<br>and signaling | Information storage<br>and processing | Metabolism |
| --- | --- | --- | --- |
| Archaea top 10% | 0.228 | 0.197 | **0.575** |
| Archaea<br>whole gene families | 0.249 | 0.246 | 0.505 |
| Micrococcales<br>top 10% | 0.241 | 0.147 | **0.613** |
| Micrococcales<br>whole gene families | 0.306 | 0.213 | 0.481 |
| Fungi top 10% | 0.348 | 0.198 | **0.455** |
| Fungi<br>whole gene families | 0.429 | 0.288 | 0.282 |

**Table 4.** Enrichment of COG categories in the metabolism supercategory by frequent gene gain/loss events.

| COG<br>category | C | E | F | G | H | I | P | Q |
| --- | --- | --- | --- | --- | --- | --- | --- | --- |
| Archaea top<br>10% | **0.15** | 0.097 | 0.016 | 0.059 | 0.048 | 0.024 | **0.138** | 0.028 |
| Archaea<br>whole gene<br>families | 0.1 | 0.095 | 0.032 | 0.076 | 0.057 | 0.031 | 0.075 | 0.030 |
| Micrococcales<br>top 10% | 0.060 | **0.124** | **0.024** | **0.153** | 0.032 | 0.031 | **0.133** | **0.034** |
| Micrococcales<br>whole gene<br>families | 0.070 | 0.092 | 0.020 | 0.097 | 0.046 | 0.030 | 0.084 | 0.030 |
| Fungi top<br>10% | **0.053** | **0.071** | **0.027** | **0.104** | **0.022** | **0.054** | **0.057** | **0.033** |
| Fungi<br>whole gene<br>families | 0.041 | 0.038 | 0.012 | 0.069 | 0.017 | 0.031 | 0.048 | 0.014 |

COG category symbols: C, energy production and conversion; E, amino acid transport and metabolism; F, nucleotide transport and metabolism; G, carbohydrate transport and metabolism; H, coenzyme transport and metabolism; I, lipid transport and metabolism; P, inorganic ion transport and metabolism; Q, secondary metabolites biosynthesis, transport, and catabolism.

## 4. Discussion

In this study, we developed Mirage, which adopts the RER model and phylogenetic mixture model for accurate ML reconstruction of gene-content evolutionary history. We demonstrated that the combination achieved good performance based on simulated and empirical datasets.

The differences between the RER model and the UNREST model in DNA evolution models are noteworthy, because elements of the UNREST evolutionary rate matrix are independent of each other (Yang 1994). The first difference lies in the way of estimating *π*. Whereas *π* is modeled as the stationary distribution of the Markov process formulated by the parameter matrix 3 in the UNREST model, *π* and *R* are defined as independent parameters in the RER model, because of the difficulty of assuming the stationarity in the gene-content evolution. Second, the RER model assumes that [*R*]_*i,j*_ = 0 when |*i* − *j*| > 1, like all other gene-content evolution models. Although simultaneous acquisitions/ losses of multiple genes may occur during evolution, they can be modeled as consecutive gene acquisitions/losses in a very small period of time by the RER model.

The setting of the gene-content category number *K* is also an important problem in Mirage. Although the Akaike Information Criterion (AIC) is a widely used estimator for model selection in phylogenetics, AIC can only be applied to statistically regular models, and the phylogenetic mixture model is a non-regular model. Another popular technique for model selection is the non-parametric Bayesian method. This method can be applied to non-regular models, but requires a lot of computation time. A practical model selection method for non-regular models is an unsolved problem in statistics, and various methods have been proposed (Fujimaki and Morinaga 2012; Watanabe 2013). The integration of Mirage with model selection methods for non-regular models is a future task.

As an application of Mirage, we envision function prediction of function-unknown genes by integrating it with phylogenetic profiling to be an interesting direction. The phylogenetic profiling method predicts gene functions based on correlated occurrence patterns between genes in an ortholog table (Kensche *et al*. 2008; Kumagai *et al*. 2018; Sherill-Rofe *et al*. 2019). The method generally ignores evolutionary relationships, for example by using simple mutual information as an index of correlation, and this ignorance is known to decrease prediction performance (Kensche *et al*. 2008). Previous studies have shown that the prediction performance can be improved by observing correlation patterns of gene gain/loss events in the reconstructed gene-content evolutionary history instead of gene occurrence patterns in extant species (Barker *et al*. 2007; Ta *et al*. 2011; Moi *et al*. 2020). Thus, precise reconstruction of the gene-content evolutionary history by Mirage would contribute to the improvement of the phylogenetic profiling method.

With the present Mirage implementation, it is still difficult to reconstruct the gene-content evolutionary histories of all genome-sequenced species at once because of the huge computation time required. Thus, improving the computation time of Mirage is essential. In particular, application of the series acceleration method, which improves the convergence rate of a series, to the iteration steps of the EM algorithm seems promising. Specifically, the vector-ɛ acceleration technique, which does not require derivation of acceleration formula for each statistical model, may be readily applied to Mirage (Kuroda and Sakakihara. 2006). Another powerful approach would be a partitioning method, which does not use probabilistic but deterministic assignment of gene families to each gene cluster in the phylogenetic mixture model. This method has been widely used in molecular evolutionary analyses, but not in gene-content evolutionary analyses (Brown and Lemmon. 2007; Frandsen *et al*. 2015; Lanfear *et al*. 2017). Although the partitioning method can be less accurate due to the deterministic approximation, its computational efficiency would be high.

Although Mirage can model differences in the evolutionary rates among gene-content clusters, it assumes the same evolutionary rate among all branches of the phylogenetic tree. However, this assumption does not always hold true. For example, polyploidization events cause massive gene gains (Inoue *et al*. 2015; Sriswasdi *et al*. 2016), and parasitization events cause massive gene losses (Sun *et al*. 2018). Moreover, heterogeneity of evolutionary rates among branches may also be caused by changes in survival strategies (Sriswasdi *et al*. 2017) or large-scale extinction events (Wolf and Koonin 2013). Our knowledge of the nature of branch-wide gene gain/loss rate heterogeneities is still very limited, and the expansion of Mirage in this direction would be needed to deepen our understanding. Iwasaki and Takagi developed a heterogeneous gene-content evolution model, but their model had a problem with an increase in parameter numbers by assigning independent parameters for each branch (Iwasaki and Takagi 2007). Instead, a branch-level parameter mixture model that divides *branches* into clusters with the same evolutionary model parameters may be promising.

## 5. Materials and Methods

### 5.1 Preparation of simulated datasets

To prepare simulated datasets, we used a perfect binary tree with 128 leaves as the input phylogenetic tree topology and determined the branch lengths by randomly sampling from a uniform distribution between 0.01 and 0.05. We set the number of gene-content clusters L to 4 and the maximum gene family size *l*_*max*_ to 3. The parameter *φ* was set to (0.5, 0.2, 0.2, 0.1)^T^. We defined *r*_*i*_ as a vector ([*R*_*i*_]_0,1_, [*R*_*i*_]_1,2_, [*R*_*i*_]_2,3_)^T^ for simplicity of notation, and we set *r*_1_, *r*_2_, *r*_3_, and *r*_4_ to (0.05, 0.05, 0.05)^T^, (0.2, 0.3, 0.5)^T^, (2.0, 0.5, 0.5)^T^, and (2.5, 3.0, 4.5)^T^, respectively. We used the same values for [*R*_*i*_]_*j*,*j*+1_ and [*R*_*i*_]_*j*,*j*−1_ for any *j* Intuitively, the four clusters represent: 1) a cluster with few gene-copy number changes, 2) a cluster with moderate changes, 3) a cluster with a large variation between 0 and 1, and 4) a cluster with large overall variations. Finally, we set *π*_1_, *π*_2_, *π*_3_, and *π*_4_ to (0.8, 0.15, 0.04, 0.01)^T^, (0.1, 0.7, 0.1, 0.1)^T^, (0.45, 0.45, 0.05, 0.05)^T^, and (0.4, 0.1, 0.1, 0.4)^T^, respectively.

We simulated the evolution of 10,000 gene families along the input phylogenetic tree for a simulated dataset, and we constructed an ortholog table from the gene copy numbers at the leaf nodes. We prepared 10 simulated datasets, each of which consisted of an ortholog table and a phylogenetic tree.

### 5.2 Parameter estimation and the data analysis

The EM algorithm is guaranteed to converge to a local optimum but not to a global optimum, and thus, the estimation results can depend on the initial values of the model parameters. Therefore, we estimated the parameters 100 times using the EM algorithm for each dataset and each model setting, and we adopted the estimation results with the highest data likelihood in the training dataset.

We compared the RER model with the two-parameter, C&M, and BDI models with restriction of the maximum value of the gene copy number value. Because a software program to estimate the parameters under these model settings was not available, we implemented our own software. Technical details are described in the Supplementary Materials. In the simulated dataset analysis, we used model settings that mimic those in previous studies for comparison. Specifically, we investigated the model settings of (1) the RER model (*K* = 4, *l*_*max*_ = 1) (Cohen and Pupko 2010), (2) the two-parameter model (*K* = 1, *l*_*max*_ = 3) (Hahn *et al*. 2005; Iwasaki and Takagi 2007), (3) the C&M model (*K* = 1, *l*_*max*_ = 3) (Csurös and Miklós 2009) and (4) the BDI model (*K* = 1, *l*_*max*_ = 3) (Karev *et al*. 2002; Ames *et al*. 2012). Additionally, we investigated the model setting that changed L of the true model setting from 4 to 1 to assess the effect of the phylogenetic mixture model on the performance.

As the evaluation criteria for the reconstructed evolutionary history in the simulated dataset analysis, we used the proportion of gene families whose gene copy numbers were correctly estimated in ancestral nodes, except for the two-state model. For the two-state model and the true model settings, we evaluated the estimation accuracy of the presence or absence of gene families. We used the averaged accuracy from the 10 simulated datasets.

In the empirical dataset analysis, we tested *K* values from 1 to 15 and *l*_*max*_ values from 2 to 4. We investigated the two-parameter, C&M, BDI model, and RER models as evolutionary models.

### 5.3 Preprocessing of the empirical datasets

We extracted gene family data of Archaea, Micrococcales, and Fungi genomes from the ortholog table in the STRING database version 11.0 (Szklarczyk *et al*. 2019). We used the NCBI Taxonomy database for taxonomic annotation. Next, we retrieved species in the phylogenetic trees provided by the Genome Taxonomy Database release 89 (Parks *et al*. 2019) for Archaea and Micrococcales, and those provided by the SILVA database release 111 (Yilmaz *et al*. 2014; Yarza *et al*. 2017) for Fungi. Then, we removed species data that were only included in either the ortholog tables or the phylogenetic trees from these datasets. In the Fungi phylogenetic tree, for some species, there were multiple strains per species. For these species, we randomly selected one strain and removed the others. Then, we reshaped the phylogenetic trees to satisfy the following three conditions: (1) a leaf of the phylogenetic trees always corresponds to a species, (2) tree topology of the tree is a binary tree, and (3) the distances and the phylogenetic relationships between species are the same as in the original tree. Because there were branches with branch lengths of 0 in the Fungi phylogenetic tree, we added a pseudo length 0.0001 to all tree branches. Note that the minimum branch length excluding 0 in the tree was 0.00059, which is larger than 0.0001. Finally, we removed their gene families shared by *N* or *N* − 1 or only one organism from the ortholog tables. The constructed datasets are freely available at https://github.com/fukunagatsu/Mirage.

### 5.4 Division of the datasets into the training and test datasets

We divided the datasets in the following three ways. For experiment 1, we randomly divided the gene families into training and test datasets at a 3:1 ratio for each dataset. In this method, the species were common between the two datasets. For experiment 2, for each dataset, we randomly divided the species into training and test datasets at a 1:1 ratio and removed the gene families that were not present or shared by only 1 or *n* − 1 or. species in each dataset after the division. Here, *n* is defined as the number of species in the dataset after the division. In this method, some gene families were shared between the training and test datasets. For experiment 3, we further processed the datasets obtained by the second method. In particular, we randomly assigned gene families shared between the training and test datasets to either of the datasets, so that no gene families were shared between the two datasets. Therefore, in this division, both the species and the gene families differed between the two datasets. The numbers of gene families for each dataset in the experiments 2 and 3 are listed in Table S7.

## Acknowledgements

This work was supported by the Japan Society for the Promotion of Science [grant numbers JP19K20395 and JP20H05582 to T.F. and 16H06279 and 19H05688 to W.I.]. Computations in this research were performed using the supercomputing facilities at the National Institute of Genetics in Research Organization of Information and Systems.

## Notes

### Competing Interest Statement

The authors have declared no competing interest.

## References

Ames RM, Money D, Ghatge VP, Whelan S, Lovell SC. 2012. Determining the evolutionary history of gene families. Bioinformatics. 28(1):48–55

Barker D, Maede A, Pagel M. 2007. Constrained models of evolution lead to improved prediction of functional linkage from correlated gain and loss of genes. Bioinformatics 23(1):14–20

Brown JM, Lemmon AR. 2007. The importance of data partitioning and the utility of Bayes factors in Bayesian phylogenetics. Syst. Biol. 56(4):643–655

Cohen O, Pupko T. 2010. Inference and characterization of horizontally transferred gene families using stochastic mapping. Mol. Biol. Evol. 27(3):703–713

Cohen O, Pupko T. 2011. Inference of gain and loss events from phyletic patterns using stochastic mapping and maximum parsimony--a simulation study. Genome Biol. Evol. 3:1265–1275

Csurös M, Miklós I. 2009. Streamlining and large ancestral genomes in Archaea inferred with a phylogenetic birth-and-death model. Mol. Biol. Evol. 26(9):2087–2095

Dang T, Kishino H. 2019. Stochastic variational inference for Bayesian phylogenetics: A case of CAT model. Mol. Biol. Evol. 36(4):825–833

Dempster AP, Nan ML, Donald BR. 1977. Maximum likelihood from incomplete data via the EM algorithm, J. R. Stat. Soc. Sect. B 39(1):1–22.

Fernández R, Gabaldón T. 2020. Gene gain and loss across the metazoan tree of life. Nat. Ecol. Evol. 4(4):524–533

Fitzpatrick DA. 2012. Horizontal gene transfer in Fungi. FEMS Microbiol. Lett. 329(1):1–8

Frandsen BP, Calcott B, Mayer C, Lanfer R. 2015. Automatic selection of partitioning schemes for phylogenetic analyses using iterative K-means clustering of site rates. BMC Evol. Biol. 15(1):13

Fujimaki R, Morinaga S. 2012. Factorized asymptotic Bayesian inference for mixture modeling. Artificial Intelligence and Statistics. 400–408

Hahn MW, De Bie T, Stajich JE, Nguyen C, Cristianini N. 2005. Estimating the tempo and mode of gene family evolution from comparative genomic data. Genome Res. 15(8):1153–1160

Holmes I, Rubin GM. 2002. An expectation maximization algorithm for training hidden substitution models. J. Mol. Biol. 317:753–764

Inoue J, Sato Y, Sinclair R, Tsukamoto K, Nishida M. 2015. Rapid genome reshaping by multiple-gene loss after whole-genome duplication in teleost fish suggested by mathematical modeling. Proc. Natl. Acad. U. S. A. 112(48):14918–14923

Iwasaki W, Takagi T. 2007. Reconstruction of highly heterogeneous gene-content evolution across the three domains of life. Bioinformatics 23(13):i230–i239

Iwasaki W, Takagi T. 2009. Rapid pathway evolution facilitated by horizontal gene transfers across prokaryotic lineages. PLoS Genet. 5(3):e1000402

Jeltsch A, Pingoud A. 1996. Horizontal gene transfer contributes to the wide distribution and evolution of type II restriction-modification systems. J. Mol. Evol. 42(2):91–96

Karev GP, Wolf YI, Rzhetsky AY, Berezovskaya FS, Koonin EV. 2002. Birth and death of protein domains: A simple model of evolution explains power law behavior. BMC Evol. Biol. 2:18

Kensche RP, van Noort V, Huynen AM. 2008. Practical and theoretical advances in predicting the function of a protein by its phylogenetic distribution. J. R. Soc. Interface. 5(19):151–170

Kiryu H. 2011. Sufficient statistics and expectation maximization algorithms in phylogenetic tree models. Bioinformatics 27(17): 2346–2353

Kumagai Y, Yoshizawa S, Nakajima Y, Watanabe M, Fukunaga T, Ogura Y, Hayashi T, Oshima K, Hattori M, Ikeuchi M, et al. 2018. Solar-panel and parasol strategies shape the proteorhodopsin distribution pattern in marine Flavobacteriia. ISME J. 12(5):1329–1343

Kuroda M, Sakakihara M. 2006. Accelerating the convergence of the EM algorithm using the vector *ε* algorithm. Comput. Stat. Data Anal. 51(3): 1549–1561

Lartillot N, Philippe H. 2004. A Bayesian mixture model for across-site heterogeneities in the amino-acid replacement process. Mol. Biol. Evol. 21(6):1095–1109

Lanfear R, Frandsen PB, Wright AM, Senfeld T, Calcott B. 2017. PartitionFinder 2: New methods for selecting partitioned models of evolution for molecular and morphological phylogenetic analyses. Mol. Biol. Evol. 34(3):772–773

Li Y, Calvo SE, Gutman R, Liu JS, Mootha VK. 2014. Expansion of biological pathways based on evolutionary inference. Cell. 158(1): 213–225.

Li Y, Ning S, Calvo SE, Mootha VK, Liu JS. 2019. Bayesian hidden Markov tree models for clustering genes with shared evolutionary history. Ann. Appl. Stat. 13(1): 606–637.

Moi D, Kilchoer L, Aguilar SP, Dessimoz C. 2020. Scalable phylogenetic profiling using MinHash uncovers likely eukaryotic sexual reproduction genes. PLoS Comput Biol.,16(7): e1007553

Montague JM, Li G, Gandolfi B, Khan R, Aken LB, Searle MJS, Minx P, Hiller WL, Koboldt DC, Davis WB et al. 2014. Comparative analysis of the domestic cat genome reveals genetic signatures underlying feline biology and domestication. Proc. Natl. Acad. U. S. A. 111(48): 17230–17235

Niimura Y. 2009. Evolutionary dynamics of olfactory receptor genes in chordates: Interaction between environments and genomic contents. Hum Genomics. 4(2):107–118

Pagel M, Meade A. 2004. A phylogenetic mixture model for detecting pattern-heterogeneity in gene sequence or character-state data. Syst. Biol. 53(4):571–581

Parks HD, Chuvochina M, Waite WD, Rinke C, Skarshewski A, Chaumeil PA, Hugenholtz P. 2018. A standardized bacterial taxonomy based on genome phylogeny substantially revises the tree of life. Nat. Biotechnol. 36(10):996–1004

Powell S, Szklarczyk D, Trachana K, Roth A, Kuhn M, Muller J, Arnold R, Rattei T, Letunic I, Doerks T et al. 2012. eggNOG v3.0: Orthologous groups covering 1133 organisms at 41 different taxonomic ranges. Nucleic Acids Res. 42(Database issue):D284–D289

Powell S, Forslund K, Szklarczyk D, Trachana K, Roth A, Huerta-Cepas J, Gabaldón T, Rattei T, Creevey C, Kuhn M et al. 2014. eggNOG v4.0: Nested orthology inference across 3686 organisms. Nucleic Acids Res. 42(Database issue):D231–D239

Pupko T, Pe’er I, Shamir R, Graur D. 2000. A fast algorithm for joint reconstruction of ancestral amino acid sequences. Mol. Biol. Evol. 17(6):890–896

Quang le S, Gascuel O, Lartillot N. 2008. Empirical profile mixture models for phylogenetic reconstruction. Bioinformatics. 24(20):2317–2323

Saitou M, Gokcumen O. 2020. An evolutionary perspective on the impact of genomic copy number variation on human health. J. Mol. Evol. 88(1):104–119

Sherill-Rofe D, Rahat D, Findlay S, Mellul A, Guberman I, Braun M, Bloch I, Lalezari A, Samiei A, Sadreyev R, et al. 2019 Mapping global and local coevolution across 600 species to identify novel homologous recombination repair genes Genome Res. 29(3):439–448

Snel B, Bork P, Huynen MA. 2002. Genomes in flux: the evolution of archaeal and proteobacterial gene content. Genome Res. 12(1):17–25

Spencer M, Sangaralingam A. 2009. A phylogenetic mixture model for gene family loss in parasitic bacteria. Mol. Biol. Evol. 26(8):1901–1908

Sriswasdi S, Takashima M, Manabe R, Ohkuma M, Sugita T, Iwasaki W. 2016. Global deceleration of gene evolution following recent genome hybridizations in Fungi. Genome Res. 26(8): 1081–1090.

Sriswasdi S, Yang CC, Iwasaki W. 2017. Generalist species drive microbial dispersion and evolution. Nat. Commun. 8(1):1162

Sun G, Xu Y, Liu H, Sun T, Zhang J, Hettenhausen C, Shen G, Qi J, Qin Y, Li J et al. 2018. Large-scale gene losses underlie the genome evolution of parasitic plant *Cuscuta Australis*. Nat. Commun. 9(1):2683

Szklarczyk D, Gable LA, Lyon D, Junge A, Wyder S, Huerta-Cepas J, Simonovic M, Doncheva TN, Morris HJ, Bork P et al. 2019. STRING v11: Protein-protein association networks with increased coverage, supporting functional discovery in genome-wide experimental datasets. Nucleic Acids Res. 47(D1):D607–D613

Ta XH, Koskinen P, Holm L. 2011. A novel method for assigning functional linkages to proteins using enhanced phylogenetic trees. Bioinformatics 27(5):700–706

Watanabe S. 2013. A widely applicable Bayesian information criterion. Journal of Machine Learning Research. 14:867–897

Wolf IY, Koonin EV. 2013. Genome reduction as the dominant mode of evolution. Bioessays. 35(9):829–837

Yang Z. 2014. Estimating the patterns of nucleotide substitution. J. Mol. Evol. 39:105–111

Yarza P, Yilmaz P, Panzer K, Glöckner FO, Reich M. 2017. A phylogenetic framework for the kingdom Fungi based on 18S rRNA gene sequences. Mar. Genomics 36:33–39

Yilmaz P, Parfrey WL, Yarza P, Gerken J, Pruesse E, Quast C, Schwer T, Peplies J, Ludwig W, Glöckner FO. 2014. The SILVA and “All-species Living Tree Project (LTP)” taxonomic frameworks. Nucleic Acids Res. 42(Database issue):D643–648

Zamani-Dahaj SA, Okasha M, Kosakowski J, Higgs PG. 2016. Estimating the frequency of horizontal gene transfer using phylogenetic models of gene gain and loss. Mol. Biol. Evol. 33(7):1843–1857

