## Supplementary Materials for "Mirage: A phylogenetic mixture model to reconstruct gene-content evolutionary history using a realistic evolutionary rate model"

#### Supplementary Text

##### Details of Mirage's EM algorithm

The Q function of our EM algorithm is described as follows:

$$Q(\theta, \theta^{old}) = \frac{1}{L}\sum_{l=1}^{L}\sum_{k=1}^{K}\sum_{X\in\Omega(D_l)} p(X, Z_{lk}|D_l, \theta^{old}) \ln p(X, Z_{lk}, D_l|\theta)$$

, where $D_l$ is the column $l$ of the ortholog table $D$, $\Omega(D_l)$ is a set of all possible gene content evolutionary histories on the $D_l$, and $Z_{lk}$ is an indicator variable representing whether the gene family $l$ belongs to the gene content cluster $k$. Here,

$$\begin{aligned}\ln p(X, Z_{lk}, D_l|\theta) &= \ln p(X, Z_{lk}|\theta)\\ &= \ln p(X|Z_{lk}, \theta) + \ln p(Z_{lk}|\theta)\\ &= \ln p(X|R_k, \pi_k) + \ln \phi_k.\end{aligned}$$

Therefore,

$$\begin{aligned}Q(\theta, \theta^{old}) &= \frac{1}{L}\sum_{l=1}^{L}\sum_{k=1}^{K} p(Z_{lk}|D_l, \theta^{old}) \sum_{X\in\Omega(D_l)} p(X|R_k^{old}, \pi_k^{old})(\ln \phi_k + \ln p(X|R_k, \pi_k))\\ &= \frac{1}{L}\sum_{l=1}^{L}\sum_{k=1}^{K} p(Z_{lk}|D_l, \theta^{old})\Big(\ln \phi_k + \sum_{X\in\Omega(D_l)} p(X|R_k^{old}, \pi_k^{old}) \ln p(X|R_k, \pi_k)\Big).\end{aligned}$$

Here,

$$\begin{aligned}p(Z_{lk}|D_l, \theta) &\propto p(D_l|Z_{lk}, \theta)p(Z_{lk}|\theta) = \phi_k p(D_l|R_k, \pi_k)\\ \therefore \gamma(Z_{lk}) &\coloneqq p(Z_{lk}|D_l, \theta) = \frac{\phi_k p(D_l|R_k, \pi_k)}{\sum_{j=1}^{K} \phi_j p(D_l|R_j, \pi_j)}.\end{aligned}$$

Besides, by a discussion of the sufficient statistics for the phylogenetic tree model [1],

$$p(X|R_k, \pi_k) = \sum_{m=1}^{M}\sum_{i=0}^{l_{max}} t_m[R_k]_{ii} F^{(m)}(i, X) + \sum_{m=1}^{M}\sum_{i=0}^{l_{max}}\sum_{j=0, j\neq i}^{l_{max}} \ln(t_m[R_k]_{ij}) N^{(m)}(i, j, X) + \sum_{i=0}^{l_{max}} n^{root}(i, X)\ln(\pi_{ki})$$

, where $m$ is the index of node, $M$ is the number of nodes other than the root node, $t_m$ is the length of a branch between the node $m$ and the parent node. $F^{(m)}(i, X)$ and $N^{(m)}(i, j, X)$ are the fractional durations of state $i$ and the number of state changes from $i$ to $j$ at the branch between the node $m$ and the parent node on the history $X$, respectively. $n^{root}(i, X)$ is an indicator variable representing whether the root node takes the state $i$ on the history $X$. By substituting these formulas into the Q function, we obtained the following equation:

$$Q(\theta, \theta^{old}) = \frac{1}{L}\sum_{l=1}^{L}\sum_{k=1}^{K}\gamma(Z_{lk})\Bigg(\ln\phi_k + \sum_{m,i} t_m[R_k]_{ii}F^{(m)}(i, D_l, R_k^{old}, \pi_k^{old}) + \sum_{m,i,j}\ln(t_m[R_k]_{ij})N^{(m)}(i, j, D_l, R_k^{old}, \pi_k^{old}) + \sum_i n^{root}(i, D_l, R_k^{old}, \pi_k^{old})\ln(\pi_{ki})\Bigg)$$

, where $F^{(m)}(i, D_l, R_k^{old}, \pi_k^{old})$, $N^{(m)}(i, j, D_l, R_k^{old}, \pi_k^{old})$, and $n^{root}(i, D_l, R_k^{old}, \pi_k^{old})$ are the expected values of $F^{(m)}(i, X)$, $N^{(m)}(i, j, X)$, and $n^{root}(i, X)$ given $D_l$, $R_k^{old}$ and $\pi_k^{old}$, respectively.

In the step 2 of our EM algorithm, we calculated the values of $\gamma(Z_{lk})$, $F^{(m)}(i, D_l, R_k^{old}, \pi_k^{old})$, $N^{(m)}(i, j, D_l, R_k^{old}, \pi_k^{old})$, and $n^{root}(i, D_l, R_k^{old}, \pi_k^{old})$ for each $k$ and $l$. These expected values can be efficiently calculated using eigenvalue decompositions of the state transition probability matrices and a dynamic programming method for a phylogenetic tree [1].

Next, we found the parameter $\theta$ that maximized the Q function in the step 3. The calculation method is as follows. We define $\alpha_{k,i}$ and $\beta_{k,i}$ as $[R_k]_{i,i+1}$ and $[R_k]_{i,i-1}$, respectively. Then,

$$[R_k]_{i,i} = \begin{cases} -\alpha_{k,0} & (i = 0) \\ -\alpha_{k,i} - \beta_{k,i} & (1 \leq i < l_{max}) \\ -\beta_{k,l_{max}} & (i = l_{max}) \end{cases} .$$

Therefore, the calculations of $\alpha_{k,i}$ and $\beta_{k,i}$ are

$$\frac{\partial Q}{\partial \alpha_{k,i}} = 0 \Leftrightarrow \alpha_{k,i} = \frac{\sum_{l=1}^{L}\gamma(Z_{lk})\sum_m N^{(m)}(i, i+1, D_l, R_k, \pi_k)}{\sum_{l=1}^{L}\gamma(Z_{lk})\sum_m t_m F^{(m)}(i, D_l, R_k, \pi_k)} \text{ and}$$

$$\frac{\partial Q}{\partial \beta_{k,i}} = 0 \Leftrightarrow \beta_{k,i} = \frac{\sum_{l=1}^{L}\gamma(Z_{lk})\sum_m N^{(m)}(i, i-1, D_l, R_k, \pi_k)}{\sum_{l=1}^{L}\gamma(Z_{lk})\sum_m t_m F^{(m)}(i, D_l, R_k, \pi_k)}.$$

In addition, $\pi_{k,i}$ and $\phi_i$ are calculated as follows using the method of Lagrange multiplier:

$$\frac{\partial Q}{\partial \pi_{k,i}} = 0 \Leftrightarrow \pi_{k,i} = \frac{\sum_{l=1}^{L}\gamma(Z_{lk})n^{root}(i, D_l, R_k^{old}, \pi_k^{old})}{\sum_{l=1}^{L}\sum_{k=1}^{K}\gamma(Z_{lk})n^{root}(i, D_l, R_k^{old}, \pi_k^{old})} \text{ and}$$

$$\frac{\partial Q}{\partial \phi_k} = 0 \Leftrightarrow \phi_k = \frac{\sum_{l=1}^{L}\gamma(Z_{lk})}{\sum_{l=1}^{L}\sum_{k=1}^{K}\gamma(Z_{lk})}.$$

In the step 1, we initialized the parameters $\alpha_{k,i}$ and $\beta_{k,i}$ by randomly sampling from a uniform distribution between 0.5 and 5.0. We also randomly initialized the parameters $\pi_{k,i}$ and $\phi_k$ to satisfy the conditions of the probability distribution. We terminated the EM algorithm in the step 4 when the increase of the log likelihood was less than 1.0 compared to the previous iteration or the number of the iteration of the step 2 and step 3 exceeded 200.

#### An algorithm for reconstruction of gene content evolutionary history

We reconstructed the gene content evolutionary history based on a method roughly similar to the one presented by Pupko *et al.* [2]. For a gene family, we define $L_x(i,k)$ as the likelihood of the maximum likelihood reconstruction of the subtree rooted at a node $x$ when the state of the parent node of $x$ is $i$ and the gene family belongs to a gene content cluster $k$. We also define $C_x(i,k)$ as the state of the node $x$ in that reconstruction. We do not define these values for the root node. We calculated these values from the leaf nodes to the root node by a dynamic programming method. The detail is as follows:

1. Let $a$ be the state at a leaf node $x$. For each state $i$ and each gene content cluster $k$, we set $C_x(i,k)$ to $a$ and $L_x(i,k)$ to $P(a|i,R_k,t_x)$, where $t_x$ is a branch length between the node $x$ and the parent node. We perform the calculation for all leaf nodes.

2. We consider a non-root internal node $x$ whose children's $C_x(i,k)$ and $L_x(i,k)$ have already been calculated. Let $y$ and $z$ be the children of the node $x$. Here, $L_x(i,k)$ is set to $\max_j P(j|i,R_k,t_x) \times L_y(j,k) \times L_z(j,k)$ and $C_x(i,k)$ is set to $j$ achieving the maximum. We calculate these values for all non-root internal nodes based on a dynamic programming method.

3. Let $y$ and $z$ be the children of the root node. We calculate the values $\pi_k L_y(j,k) \times L_z(j,k)$ for each state $j$ and each gene content cluster $k$. We select $j$ and $k$ maximizing the value as the reconstructed state of the root node and the gene content cluster to which the gene family belongs, respectively. In the next step, the value of $k$ is fixed to the value selected in this step.

4. We consider a non-root internal node $x$ whose parent's state has already been reconstructed. When we define $i$ as the state at the parent of the node $x$, we select $C_x(i,k)$ as the reconstruction of node $x$. We reconstruct the states for all non-root internal nodes based on a dynamic programming method.

#### Parameter estimation algorithms for different evolutionary models

To compare the performances of the realistic evolutionary rate model with those of different evolutionary models, we implemented the parameter estimation algorithms for the two-parameter model, the C&M model, and the BDI model with the phylogenetic mixture model. As with the parameter estimation of the realistic evolutionary rate model with the phylogenetic mixture model, we used the EM algorithms for the parameter estimation of these models. The calculation of the step 1, 2, and 4 and the estimation of $\pi_k$ and $\phi$ in the step 3 are exactly the same as the aforementioned EM algorithm, and only the calculation of $R_k$ is different. For the two-parameter model, the calculation of $R_k$ in the step 3 is

$$\frac{\partial Q}{\partial \alpha_k} = 0 \Leftrightarrow \alpha_k = \frac{\sum_{l=1}^{L} \gamma(Z_{lk}) \sum_i \sum_m N^{(m)}(i, i+1, D_l, R_k, \pi_k)}{\sum_{l=1}^{L} \gamma(Z_{lk}) \sum_i \sum_m t_m F^{(m)}(i, D_l, R_k, \pi_k)} \text{ and}$$
$$\frac{\partial Q}{\partial \beta_k} = 0 \Leftrightarrow \beta_k = \frac{\sum_{l=1}^{L} \gamma(Z_{lk}) \sum_i \sum_m N^{(m)}(i, i-1, D_l, R_k, \pi_k)}{\sum_{l=1}^{L} \gamma(Z_{lk}) \sum_i \sum_m t_m F^{(m)}(i, D_l, R_k, \pi_k)}.$$

The parameter $\beta_k$ estimation for the C&M model is

$$\frac{\partial Q}{\partial \beta_k} = 0 \Leftrightarrow \beta_k = \frac{\sum_{l=1}^{L} \gamma(Z_{lk}) \sum_i \sum_m N^{(m)}(i, i-1, D_l, R_k, \pi_k)}{\sum_{l=1}^{L} \gamma(Z_{lk}) \sum_i \sum_m t_m i F^{(m)}(i, D_l, R_k, \pi_k)}.$$

Because we could not obtain the explicit updated formulas of $\alpha_k$ and $\gamma_k$ for the C&M model, we updated these parameters using a gradient descent method. The learning rate was initially set to 0.1. When the newly estimated parameter was less than 0.0 or the value of the Q function based on the new parameter increased, we multiplied the learning rate by 0.1 and re-estimated the parameter. Finally, when the learning rate was less than $1.0 \times 10^{-5}$, we finished the iteration of the gradient descent method. The learning rate was initialized at each iteration of the EM algorithm.

For the BDI model, the parameter estimation is

$$\frac{\partial Q}{\partial \alpha_k} = 0 \Leftrightarrow \alpha_k = \frac{\sum_{l=1}^{L} \gamma(Z_{lk}) \sum_{i,i\neq 0} \sum_m N^{(m)}(i, i+1, D_l, R_k, \pi_k)}{\sum_{l=1}^{L} \gamma(Z_{lk}) \sum_{i,i\neq 0} \sum_m t_m F^{(m)}(i, D_l, R_k, \pi_k)},$$

$$\frac{\partial Q}{\partial \beta_k} = 0 \Leftrightarrow \beta_k = \frac{\sum_{l=1}^{L} \gamma(Z_{lk}) \sum_i \sum_m N^{(m)}(i, i-1, D_l, R_k, \pi_k)}{\sum_{l=1}^{L} \gamma(Z_{lk}) \sum_i \sum_m t_m F^{(m)}(i, D_l, R_k, \pi_k)}, \text{ and}$$

$$\frac{\partial Q}{\partial \delta_k} = 0 \Leftrightarrow \delta_k = \frac{\sum_{l=1}^{L} \gamma(Z_{lk}) \sum_m N^{(m)}(0, 1, D_l, R_k, \pi_k)}{\sum_{l=1}^{L} \gamma(Z_{lk}) \sum_m t_m F^{(m)}(i, D_l, R_k, \pi_k)}.$$

### Supplementary Figures

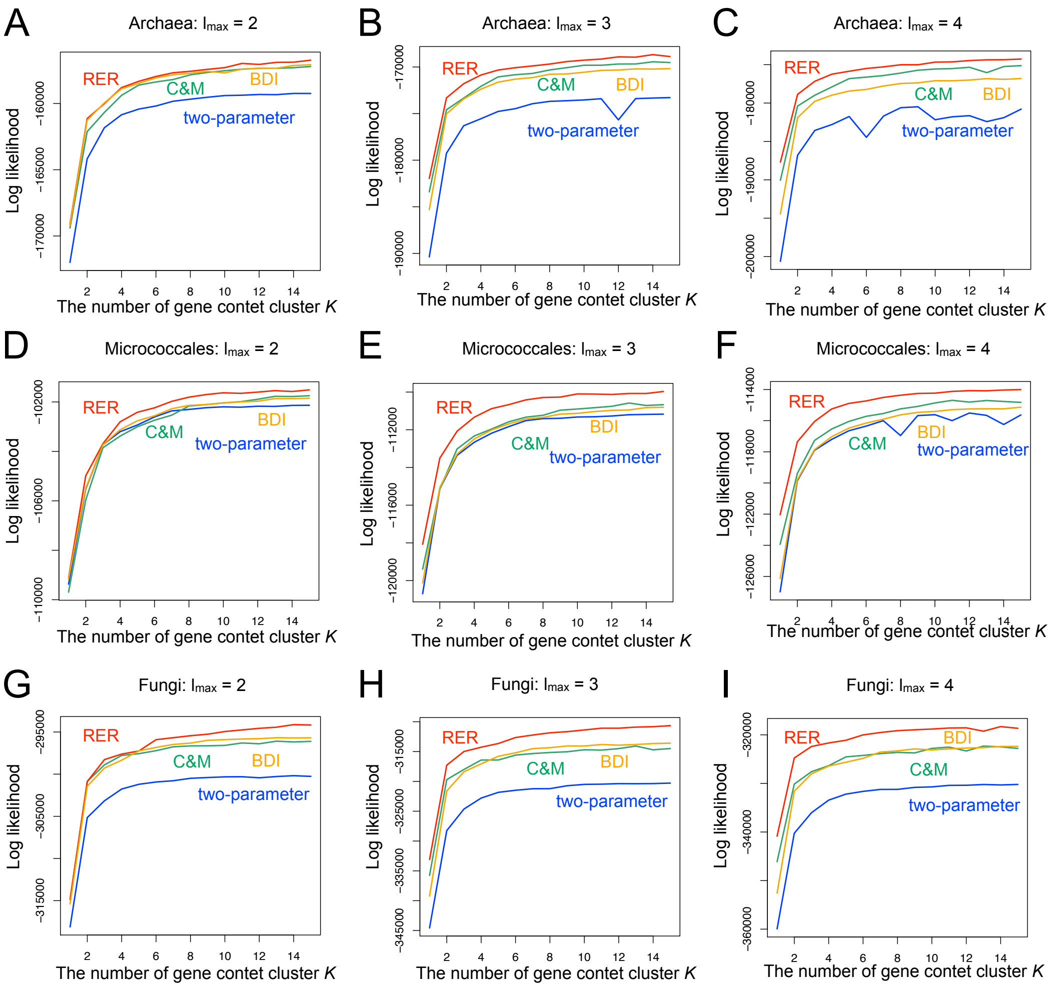


Fig. S1 The comprehensive comparison results of various model settings for the experiment 2. The x-axis and the y-axis represent the number of gene content cluster $K$ and the log-likelihood of the test dataset, respectively. The two-parameter model, the C&M model, the BDI model, and the RER model are represented by blue, green, yellow, and red lines, respectively. The panels represent the comparison results for (A-C) the Archaea dataset when $l_{max}$ was set to 2-4, (D-F) the Micrococcales dataset when $l_{max}$ was set to 2-4, and (G-I) the Fungi dataset when $l_{max}$ was set to 2-4.

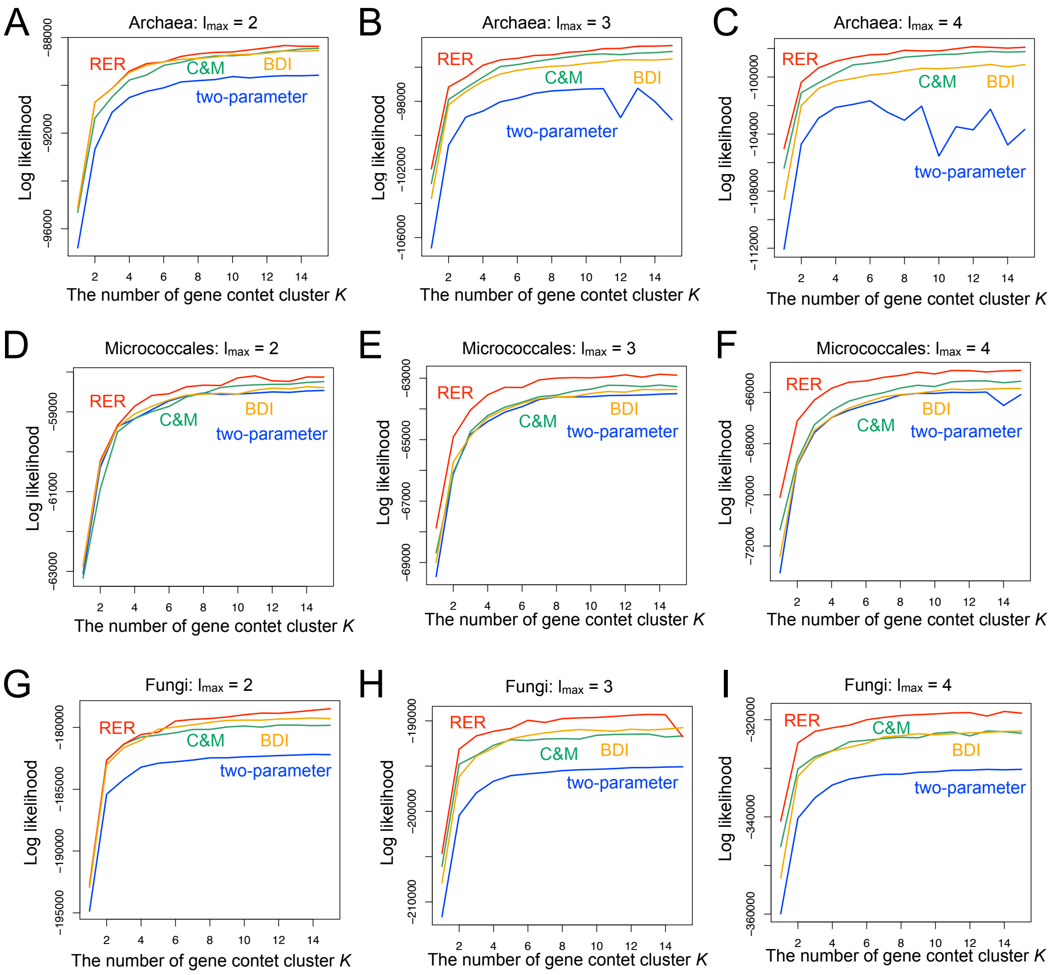


Fig. S2 The comprehensive comparison results of various model settings for the experiment 3. The x-axis and the y-axis represent the number of gene content cluster $K$ and the log-likelihood of the test dataset, respectively. The two-parameter model, the C&M model, the BDI model, and the RER model are represented by blue, green, yellow, and red lines, respectively. The panels represent the comparison results for (A-C) the Archaea dataset when $l_{max}$ was set to 2-4, (D-F) the Micrococcales dataset when $l_{max}$ was set to 2-4, and (G-I) the Fungi dataset when $l_{max}$ was set to 2-4.

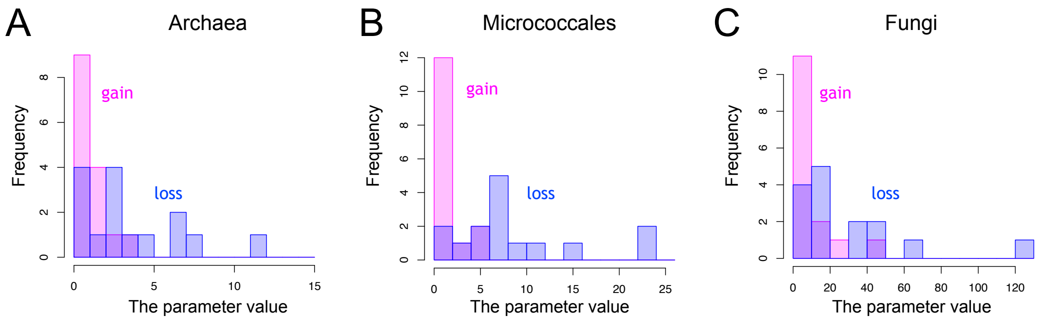


Fig. S3 The distribution of the estimated gene gain and loss rates in the empirical datasets. The x-axis and the y-axis represent the estimated parameter value and the frequency, respectively. The gene gain and loss rates are represented by red and blue bars, respectively. The panels represent the distributions for (A) the Archaea dataset, (B) the Micrococcales dataset, and (C) the Fungi dataset.

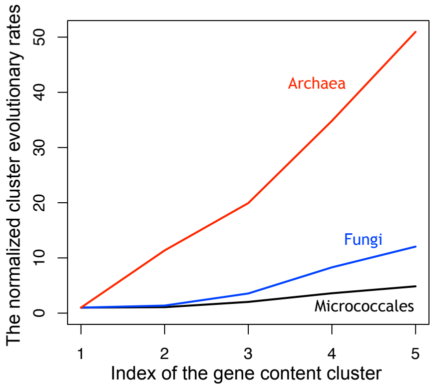


Fig. S4 The values of the normalized cluster evolutionary rates for each dataset. The x-axis and the y-axis represent the index of the gene content cluster and the normalized cluster evolutionary rates, respectively. The gene content cluster was sorted by the normalized cluster evolutionary rates. The Archaea dataset, the Micrococcales dataset, and the Fungi dataset are represented by red, black, and blue lines, respectively.

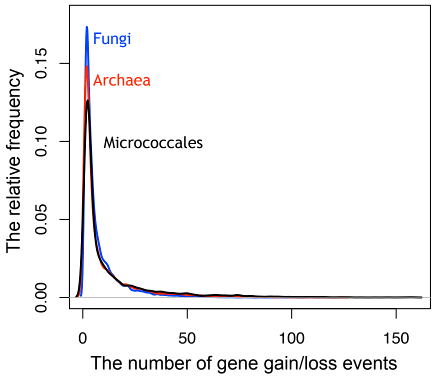


Fig. S5 The distribution of the number of gene gain/loss events for gene families in the reconstructed evolutionary history. The x-axis and the y-axis represent the number of gene gain/loss events and the relative frequency, respectively. The Archaea dataset, the Micrococcales dataset, and the Fungi dataset are represented by red, black, and blue lines, respectively.

#### Supplementary Tables

Table S1 The distribution of COG supercategory of the gene families for each gene content cluster in the Archaea dataset.

| Cluster ID | Normalized cluster evolutionary rate | Cellular process and signaling | Information storage and processing | Metabolism |
|---|---|---|---|---|
| 1 | 1.00 | 0.173 | 0.269 | 0.558 |
| 2 | 50.96 | 0.346 | 0.276 | 0.378 |
| 3 | 34.84 | 0.283 | 0.228 | 0.489 |
| 4 | 11.36 | 0.140 | 0.228 | 0.632 |
| 5 | 19.93 | 0.207 | 0.22 | 0.573 |

Table S2 The distribution of COG supercategory of the gene families for each gene content cluster in the Micrococcales dataset.

| Cluster ID | Normalized cluster evolutionary rate | Cellular process and signaling | Information storage and processing | Metabolism |
|---|---|---|---|---|
| 1 | 4.86 | 0.331 | 0.199 | 0.469 |
| 2 | 1.08 | 0.356 | 0.256 | 0.388 |
| 3 | 1.00 | 0.245 | 0.193 | 0.563 |
| 4 | 3.60 | 0.229 | 0.188 | 0.583 |
| 5 | 2.03 | 0.250 | 0.183 | 0.567 |

Table S3 The distribution of COG supercategory of the gene families for each gene content cluster in the Fungi dataset.

| Cluster ID | Normalized cluster evolutionary rate | Cellular process and signaling | Information storage and processing | Metabolism |
|---|---|---|---|---|
| 1 | 8.30 | 0.455 | 0.187 | 0.358 |
| 2 | 3.56 | 0.464 | 0.294 | 0.243 |
| 3 | 1.37 | 0.388 | 0.271 | 0.34 |
| 4 | 12.05 | 0.384 | 0.292 | 0.324 |
| 5 | 1.00 | 0.414 | 0.311 | 0.275 |

Table S4 The list of gene families with frequent gene gain/loss events for the Archaea dataset

| COG ID | gene name | frequency | function |
|---|---|---|---|
| COG4743 | *COG4743* | 124 | Uncharacterized membrane protein |
| COG1668 | *NatB* | 123 | ABC-type Na+ efflux pump |
| COG3039 | *IS5* | 116 | Transposase and inactivated derivatives |
| COG1487 | *VapC* | 114 | Predicted nucleic acid-binding protein |
| COG1848 | *COG1848* | 114 | Predicted nucleic acid-binding protein |
| COG0395 | *UgpE* | 113 | ABC-type glycerol-3-phosphate transport system |
| COG0399 | *WecE* | 112 | dTDP-4-amino-4,6-dideoxygalactose transaminase |
| COG3385 | *InsG* | 111 | IS4 transposase |
| COG1804 | *CaiB* | 111 | Crotonobetainyl-CoA |
| COG1216 | *GT2* | 108 | Glycosyltransferase |
| COG2304 | *YfbK* | 108 | Secreted protein |
| COG1672 | *AAAA* | 106 | Predicted ATPase |
| COG0677 | *WecC* | 105 | UDP-N-acetyl-D-mannosaminuronate dehydrogenase |
| COG3335 | *Transposase* | 105 | Transposase |
| COG1055 | *ArsB* | 104 | Na+/H+ antiporter NhaD |
| COG1708 | *COG1708* | 104 | Predicted nucleotidyltransferase |
| COG3677 | *InsA* | 104 | Transposase |
| COG3316 | *Rve* | 102 | Transposase |
| COG3415 | *Transposase* | 104 | Transposase |
| COG0286 | *HsdM* | 101 | Type I restriction-modification system |

Table S5 The list of gene families with frequent gene gain/loss events for the Micrococcales dataset

| COG ID | gene name | frequency | function |
|---|---|---|---|
| COG3547 | *COG3547* | 158 | Transposase |
| COG3629 | *DnrI* | 158 | DNA-binding transcriptional activator of the SARP family |
| COG3293 | *COG3293* | 156 | Transposase |
| COG3209 | *RhsA* | 155 | Uncharacterized conserved protein |
| COG3328 | *IS285* | 152 | Transposase |
| COG3464 | *COG3464* | 147 | Transposase |
| COG3119 | *AslA* | 145 | Arylsulfatase A |
| COG1484 | *DnaC* | 140 | DNA replication protein |
| COG4941 | *COG4941* | 133 | Predicted RNA polymerase sigma factor |
| COG1848 | *COG1848* | 128 | Predicted nucleic acid-binding protein |
| COG4118 | *Phd* | 128 | Antitoxin component of toxin-antitoxin stability system |
| COG5001 | *COG5001* | 126 | Predicted signal transduction protein |
| COG0286 | *HsdM* | 125 | Type I restriction-modification system |
| COG3669 | *AfuC* | 124 | Alpha-L-fucosidase |
| COG2334 | *SrkA* | 123 | Ser/Thr protein kinase RdoA |
| COG1476 | *XRE* | 121 | DNA-binding transcriptional regulator |
| COG1302 | *YloU* | 119 | Uncharacterized conserved protein |
| COG0053 | *FieF* | 117 | Divalent metal cation (Fe/Co/Zn/Cd) transporter |
| COG0402 | *SsnA* | 116 | Cytosine/adenosine deaminase |
| COG3391 | *YncE* | 116 | DNA-binding beta-propeller fold protein |

Table S6 The list of gene families with frequent gene gain/loss events for the Fungi dataset

| COG ID | gene name | frequency | function |
|---|---|---|---|
| NOG10249 | *NOG10249* | 117 | Uncharacterized protein |
| COG2801 | *Tra5* | 104 | Transposase InsO |
| KOG3105 | *KOG3105* | 101 | DNA-binding centromere protein B |
| NOG54750 | *NOG54750* | 101 | DNA directed polymerase |
| KOG1216 | *KOG1216* | 96 | von Willebrand factor |
| KOG1075 | *FOG* | 95 | Reverse transcriptase |
| NOG06394 | *NOG06394* | 93 | Nitrosoguanidine resistance protein |
| NOG259057 | *NOG259057* | 89 | Uncharacterized protein |
| NOG06621 | *NOG06621* | 86 | Uncharacterized protein |
| COG0252 | *AnsA* | 85 | L-asparaginase/archaeal Glu-tRNAGln amidotransferase subunit D |
| NOG258056 | *NOG258056* | 85 | Uncharacterized protein |
| COG0328 | *RnhA* | 84 | Ribonuclease HI |
| COG2220 | *UlaG* | 81 | L-ascorbate metabolism protein |
| KOG3386 | *KOG3386* | 80 | Copper transporter |
| NOG258061 | *NOG258061* | 80 | Uncharacterized protein |
| KOG1773 | *KOG1773* | 79 | Stress responsive protein |
| KOG4701 | *KOG4701* | 79 | Chitinase |
| NOG252561 | *NOG252561* | 79 | Uncharacterized protein |
| COG0435 | *ECM4* | 78 | Glutathionyl-hydroquinone reductase |
| NOG05829 | *NOG05829* | 78 | Cloroperoxidase |
| NOG07596 | *NOG07596* | 78 | Uncharacterized protein |

Table S7 The number of gene families for each dataset in the experiment 2 and 3

| taxonomic group | training (experiment2) | test (experiment2) | training (experiment3) | test (experiment3) |
|---|---|---|---|---|
| Archaea | 7647 | 7714 | 4743 | 4810 |
| Micrococcales | 4380 | 4380 | 2816 | 2816 |
| Fungi | 15773 | 15975 | 10206 | 10409 |

#### Supplementary Data

##### The estimated parameter for the Micrococcales dataset

$$\phi_1 = 0.317,\ \pi_1 = (0.001, 0.957, 0.000, 0.042)^T,\ R_1 = \begin{pmatrix} -0.092 & 0.092 & 0 & 0 \\ 6.714 & -6.843 & 0.129 & 0 \\ 0 & 6.238 & -7.505 & 1.267 \\ 0 & 0 & 1.610 & -1.610 \end{pmatrix}$$

$$\phi_2 = 0.306,\ \pi_2 = (0.944, 0.018, 0.000, 0.039)^T,\ R_2 = \begin{pmatrix} -0.716 & 0.716 & 0 & 0 \\ 23.918 & -25.608 & 1.690 & 0 \\ 0 & 23.699 & -26.221 & 2.522 \\ 0 & 0 & 9.024 & -9.024 \end{pmatrix}$$

$$\phi_3 = 0.174,\ \pi_3 = (0.693, 0.196, 0.000, 0.110)^T,\ R_3 = \begin{pmatrix} -0.110 & 0.110 & 0 & 0 \\ 0.414 & -0.626 & 0.212 & 0 \\ 0 & 7.391 & -8.313 & 0.922 \\ 0 & 0 & 10.527 & -10.527 \end{pmatrix}$$

$$\phi_4 = 0.111,\ \pi_4 = (0.329, 0.412, 0.075, 0.185)^T,\ R_4 = \begin{pmatrix} -0.919 & 0.919 & 0 & 0 \\ 2.108 & -2.597 & 0.489 & 0 \\ 0 & 4.245 & -5.428 & 1.183 \\ 0 & 0 & 5.389 & -5.389 \end{pmatrix}$$

$$\phi_5 = 0.092,\ \pi_5 = (0.490, 0.000, 0.000, 0.510)^T,\ R_5 = \begin{pmatrix} -1.824 & 1.824 & 0 & 0 \\ 6.220 & -10.807 & 4.587 & 0 \\ 0 & 15.081 & -19.422 & 4.361 \\ 0 & 0 & 7.866 & -7.866 \end{pmatrix}$$

##### The estimated parameter for the Fungi dataset

$$\phi_1 = 0.460,\ \pi_1 = (0.953, 0.038, 0.009, 0.000)^T,\ R_1 = \begin{pmatrix} -1.092 & 1.092 & 0 & 0 \\ 42.180 & -61.128 & 19.038 & 0 \\ 0 & 123.295 & -169.418 & 46.123 \\ 0 & 0 & 68.377 & -68.377 \end{pmatrix}$$

$$\phi_2 = 0.236,\ \pi_2 = (0.854, 0.115, 0.023, 0.009)^T,\ R_2 = \begin{pmatrix} -0.438 & 0.438 & 0 & 0 \\ 10.252 & -11.017 & 0.765 & 0 \\ 0 & 5.155 & -9.142 & 3.987 \\ 0 & 0 & 2.836 & -2.836 \end{pmatrix}$$

$$\phi_3 = 0.209,\ \pi_3 = (0.827, 0.165, 0.000, 0.008)^T,\ R_3 = \begin{pmatrix} -0.357 & 0.357 & 0 & 0 \\ 2.672 & -4.908 & 2.236 & 0 \\ 0 & 46.827 & -53.511 & 6.683 \\ 0 & 0 & 31.218 & -31.218 \end{pmatrix}$$

$$\phi_4 = 0.048,\ \pi_4 = (0.409, 0.369, 0.197, 0.025)^T,\ R_4 = \begin{pmatrix} -28.259 & 28.259 & 0 & 0 \\ 2.076 & -3.579 & 1.503 & 0 \\ 0 & 11.978 & -16.564 & 4.586 \\ 0 & 0 & 12.491 & -12.491 \end{pmatrix}$$

$$\phi_5 = 0.047,\ \pi_5 = (0.760, 0.087, 0.132, 0.020)^T,\ R_5 = \begin{pmatrix} -3.403 & 3.403 & 0 & 0 \\ 14.170 & -22.078 & 7.908 & 0 \\ 0 & 32.850 & -48.754 & 15.904 \\ 0 & 0 & 18.186 & -18.186 \end{pmatrix}$$

#### References


[1] Hisanori Kiryu. “Sufficient statistics and expectation maximization algorithms in phylogenetic tree models” Bioinformatics 27(17) (2011): 2346-2353.

[2] Tal Pupko, Itsik Pe, Ron Shamir and Dan Graur. “A fast algorithm for joint reconstruction of ancestral amino acid sequences.” Molecular Biology and Evolution 17(6) (2000): 890-896.
